# High genetic diversity and presence of genetic structure characterise the Corso-Sardinian endemics *Ruta corsica* and *Ruta lamarmorae* (Rutaceae)

**DOI:** 10.1101/337931

**Authors:** Marilena Meloni, Caterina Angela Dettori, Andrea Reid, Gianluigi Bacchetta, Laetitia Hugot, Elena Conti

## Abstract

Corsica and Sardinia form one of the ten areas with highest biodiversity in the Mediterranean and are considered one of the priority regions for conservation in Europe. In order to preserve the high levels of endemism and biological diversity at different hierarchical levels, knowledge of the evolutionary history and current genetic structure of Corso-Sardinian endemics is instrumental. Microsatellite markers were newly developed and used to study the genetic structure and taxonomic status of *Ruta corsica* and *Ruta lamarmorae*, rare endemics of Corsica and Sardinia, respectively, and previously considered a single species. Our analyses identified high levels of genetic variation within each species (*P*=0.883, *H_e_*=0.543 for *R. corsica; P*=0.972, *H_e_*=0.627 for *R. lamarmorae*). Intrinsic traits of the species (hermaphroditism, proterandry and polyploidy) and island-dependent factors (i.e. age, origin and history of the islands) might explain the detected high levels of genetic variation. We discovered differentiation between *R. corsica* and *R. lamarmorae*, and genetic structure within each species, which are consistent with the observation of low dispersal ability for both species. Our genetic results support the recent taxonomic classification of *R. corsica* and *R. lamarmorae* as separate species and suggest that they diverge at only few loci. One *R. corsica* population (SA) strongly differed from all other studied populations and appeared to be the product of hybridization between the two species in STRUCTURE analyses. Our results provide important insights for the conservation of the two rare endemics. Further genetic analyses are recommended for *R. lamarmorae* and for population SA (*R. corsica*).

## Introduction

The Mediterranean basin is characterised by high species richness (10.8 species/1000 km^2^, Médail & Quézel 1999) and considered one of the main “hot spots” of biodiversity in the world (Médail & Myers 2004, Thompson 2005). Additionally, this area is particularly rich in endemic taxa (about half of the approximately 25000 plant species native to this region are endemics), which are mainly concentrated in mountain chains and islands (Médail & Quézel 1997, 1999, Thompson 2005, Cañadas *et al*. 2014).

Corsica and Sardinia (Fig 1), the two largest islands of the Western Mediterranean basin, form one of the ten areas with highest biodiversity in the Mediterranean and are particularly rich in endemics (i.e., species exclusive to a restricted area) and sub-endemics (i.e. species occurring almost entirely in a restricted area; Médail & Quézel 1997, Thompson 2005, Blondel *et al*. 2010). The Sardinian flora consists of 2498 taxa, with about 11.61% endemic (290 species) and 14% sub-endemic (378 species) to the island (Conti *et al*. 2005, 2007; Bacchetta *et al*. 2012; Fenu *et al*. 2014). Corsica’s flora consists of 2325 taxa, of which ca. 10% are endemics (230 species; Jeanmonod & Gamisans 2013). Both islands are considered a major glacial refugium (Médail & Diadema 2009) and together host nine of the 50 most threatened plant species occurring in Mediterranean islands (de Montmollin & Strahm 2005).

**Fig. 1.**
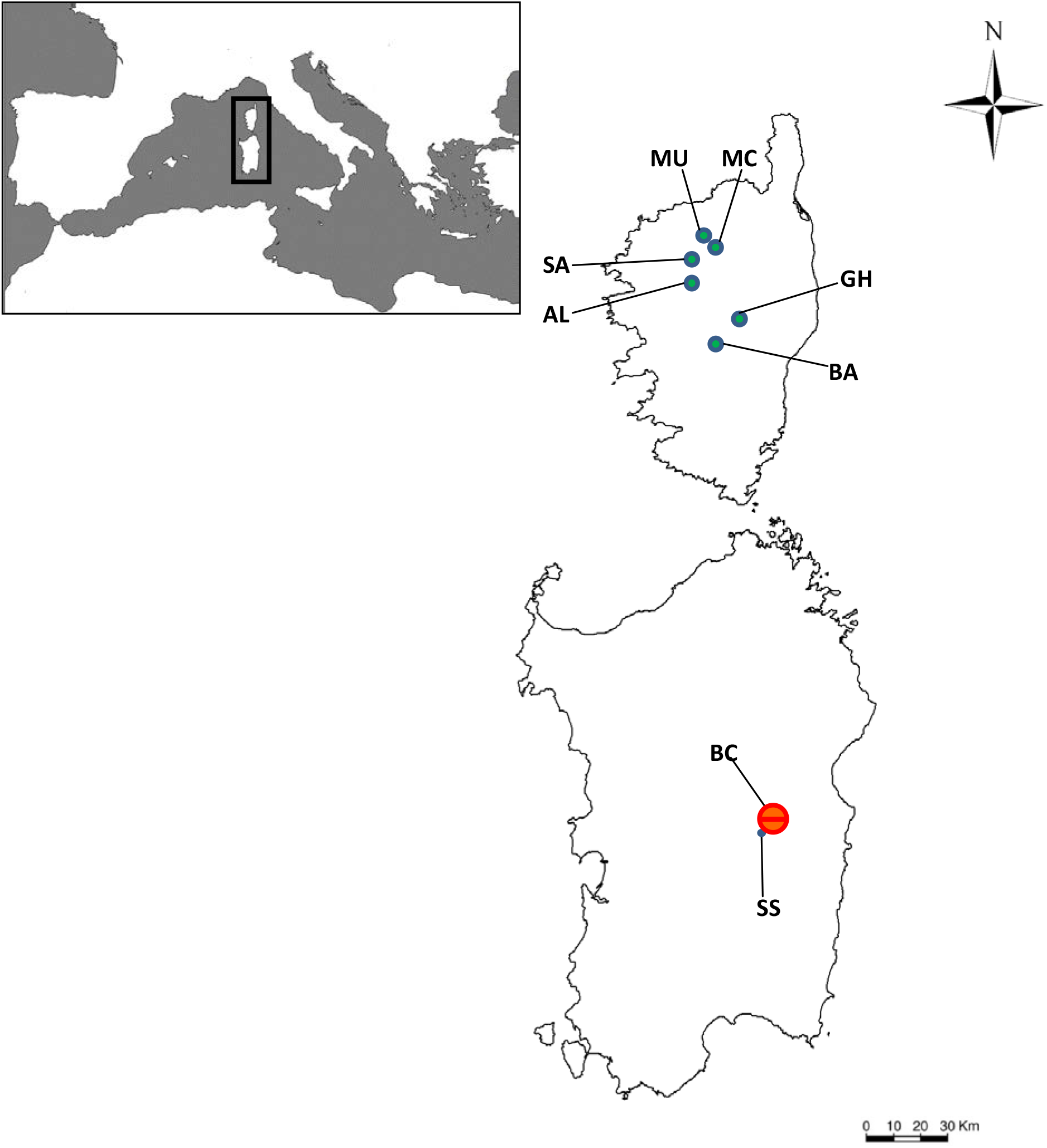
Localities of populations taxonomically assigned to *R. corsica* (green dots) and *R. lamarmorae* (red dot) sampled for this study. *Ruta lamarmorae* population was divided in two subpopulations. Detailed information on each population is provided in Table 2.

The high level of biodiversity and the number of endemics found in Corsica and Sardinia are often ascribed to their noteworthy ecosystem diversity (Bacchetta and Pontecorvo 2005) and to past geologic and paleoclimatic processes. Indeed, tectonic movements during the Tertiary (from 66 to 2.58 million years ago, MYA), the Messinian Salinity Crisis (MSC, ca. 5 MYA), the establishment of a Mediterranean climate type in the Pliocene (2-3 MYA), and climate changes associated with glaciations (Pleistocene: 0.01-2 MYA) have shaped the history of Mediterranean plant lineages (Hewitt 2000, Médail & Quézel 1997, Thompson 2005).

Corsica and Sardinia are continental fragment islands belonging to a single microplate (the Corso-Sardinian (C-S) microplate; Alvarez *et al*. 1972) and are currently separated by a narrow (11 km) and shallow (less than 50 m deep) water channel through the Bonifacio Strait. The C-S microplate was attached to Southern France and Northeastern Spain until the late Oligocene (28-30 MYA), when it broke off and rafted eastward, until it collided with the Apulian microplate (i.e., the current Italian peninsula) ca. 18-20 MYA (Rosenbaum & Lister 2004, Speranza *et al*. 2002). It reached its current position in the middle of the Western Mediterranean ca. 9 MYA (Rosenbaum & Lister 2004). The separation between Corsica and Sardinia may have begun as early as 15 MYA and was complete by 9 MYA (Alvarez 1972, 1976, Cherchi & Montadert 1982, Orsini *et al*. 1980).

Species that now occur in Corsica and Sardinia could have originated in different ways: 1) they were present in the region before the split of the C-S microplate from the Iberian peninsula; 2) they reached the C-S microplate when it was temporarily connected with the Apulian microplate during the Miocene (ca.10-20 MYA, Rosenbaum *et al*. 2002); 3) they reached the C-S microplate through the land bridges that formed among Corsica, Sardinia, the Apulian plate and the African continent during the MSC (5 MYA; Hsü *et al*. 1977, Krijgsman *et al*. 1999, McKenzie 1999) or 4) during the glacial marine regressions concomitant with the glacial cycles of the Pleistocene (Thompson 2005); 5) they reached the islands via long distance dispersal at any point in time (LDD). If insular populations differentiated sufficiently from their closest relatives due to isolation and/or extinction, they gave origin to island endemics.

Although Corsica and Sardinia are one of the priority regions for conservation in Europe (Myers *et al*. 2000, Mittermeier *et al*. 2005), knowledge of the evolution and genetic characteristics of their endemic flora, instrumental for the long-term conservation of these species, is still poor. Some molecular phylogenetic analyses have been performed to infer when and how C-S endemics reached the two islands (Yesson *et al*. 2009, Mansion *et al*. 2008, Salvo *et al*. 2008, 2010) and few studies focused on the more recent history of these species (Bacchetta *et al*. 2008; Coppi *et al*. 2008; Mameli *et al*. 2008; Bacchetta *et al*. 2011; Garrido *et al*. 2012). Nevertheless, several questions remain unanswered, including: How did C-S endemics evolve after island colonization? What is their current genetic structure? Are Corsican and Sardinian populations of the same species genetically differentiated?

*Ruta corsica* DC. and *R. lamarmorae* Bacch., Brullo & Giusso are endemics of Corsica and Sardinia, respectively. The two species belong to the small genus *Ruta*, which also includes four species widely distributed in the Mediterranean *(R. angustifolia* Pers., *R. chalepensis* L., *R. montana* L., and *R. graveolens* L.) and three species endemic to the Canary Islands (*R. oreojasme* Webb & Berth, *R. pinnata* L.f. and *R. microcarpa* Svent.). *Ruta corsica* and *R. lamarmorae* exhibit some features not found in the other species of the same genus (i.e. pulvinate, subspinescent habit; green-glaucescent leaves; white to pale yellow petals) and have been interpreted by taxonomists as relictual paleo-endemics (Bacchetta et *et al*. 2006, Cardona & Contandriopoulos 1979), in other words as ancient lineages that were more widespread in the past and are now restricted to a local region (Nekola 1999, Mishler *et al*. 2014), in this case to the C-S microplate (Arrigoni 1983, Thompson 2005). They were treated as one species (i.e., *R. corsica)* until 2006, when they were split in two different taxa based on morphological (i.e., leaf shape and size of flowers, stamens and ovaries; Bacchetta *et al*. 2006) and karyological differences *(R. corsica* is diploid, *R. lamarmorae* is tetraploid; Contandriopoulos 1957, Honsell 1957). Phylogenetic analyses of chloroplast DNA sequences from only two individuals each from Corsica and Sardinia supported the separation of *R. corsica* and *R. lamarmorae*, with individuals from the two islands grouped in mutually exclusive sister clades (Salvo *et al*. 2008).

Molecular dating analyses and inference of ancestral areas of distribution for *Ruta* species suggested that the genus originated during the Eocene in Eurasia and subsequently expanded westward and southward, colonising several landmasses of the forming Mediterranean basin (Salvo *et al*. 2010). The ancestor of the two C-S endemics likely colonised the C-S block from the Apulian plate (i.e., the emerging Italian peninsula) during the early Miocene. The divergence between the C-S endemics and the remaining *Ruta* species apparently occurred during the middle Miocene (ca. 14 MYA). Finally, *R. corsica* diverged from *R. lamarmorae* most likely in the Pliocene (ca. 3.7 MYA), when Corsica and Sardinia had already attained their current position in the middle of the Western Mediterranean sea and were separated by the Bonifacio strait. The two islands were occasionally connected by land corridors during the MSC of the Miocene and during the glacial maxima of Pleistocene climatic oscillations (Salvo *et al*. 2010).

Given the inferred biogeographic history of *R. corsica* and *R. lamarmorae* and their current distribution in Corsica and Sardinia, respectively, these species represent an ideal case study to gain new knowledge on the genetic characteristics of the Corso-Sardinian endemic flora. The aims of the present study are thus to: (1) assess the current amount and distribution of genetic diversity for the two species, testing whether the taxonomic status of *R. corsica* and *R. lamarmorae* as separate species is warranted; (2) evaluate the effects of biotic and abiotic factors (e.g., isolation, global warming, human impacts) on the genetic diversity of the two species; and (3) use the results of genetic analyses to recommend proper conservation strategies for these species, with a particular focus on *R. lamarmorae*, recently listed as endangered (Dettori *et al*. 2014a).

## Materials and methods

### Study species

*Ruta lamarmorae* was described as a species separate from *R. corsica* in 2006 (Bacchetta *et al*.). It is a small, erect, perennial shrub, 15-50 cm tall, with woody, subspinescent branches. It is characterised by bipinnate, obovate-rounded leaves, 1.5-8 cm long. The whitish, pale yellow flowers (12-13 mm in diameter) are hermaphroditic and proterandrous. The capsules, 6-7 cm long, are obtuse at the apex. As with most members of its genus, *R. lamarmorae* is tetraploid, with 2n=36 (Honsell, 1957). It blooms in June-July, fruiting in August-October. It occurs mainly on siliceous substrates, at an altitude of 1500-1750 m a.s.l. *Ruta lamarmorae* is found in a single, fragmented population in the Gennargentu massif (central-eastern Sardinia). It is categorized as endangered (EN) according to the IUCN criteria (2013; Dettori *et al*. 2014a). The main threats to this species are habitat fragmentation, overgrazing and fires (Bacchetta *et al*. 2006; Dettori *et al*. 2014a).

*Ruta corsica*, first described in 1824 by De Candolle, shares the same habit with *R. lamarmorae* and has overall similar leaves, flowers and fruits. It differs from *R. lamarmorae* in leaf shape (obovate to cuneate-oblong), smaller flower size (8-10 cm in diameter) and fruit size and shape (7-8 cm long, with apiculate apex; Bacchetta *et al*. 2006). *Ruta corsica* is the only diploid species of the genus, with 2n=18, while the other karyotyped species are tetraploid (Contandriopoulos, 1957). It blooms and fruits slightly later than *R. lamarmorae* (September-November *vs*. July-August, respectively). The species occurs at an altitude of 1000-1900 m asl and is widespread on the main Corsican massifs. It is listed as Least Concern (LC) under the French red list of threatened species (UICN, France 2013).

### Sample collection and DNA extraction

Plant material was collected during summer 2010 (see Table 2). Since the only existing population of *R. lamarmorae* has a scattered distribution on the Gennargentu massif, we sampled plants from the two opposite sides of the massif, in other words from the Northwestern (BC, 30 individuals) and Southwestern slopes (SS, 30 individuals; Fig. 1). Because sub-populations BC and SS are large (thousands of individuals; Gianluigi Bacchetta, pers. obs.), sampling was carried out in order to minimize collection of related individuals and cover the entire occupied area. Samples of *R. corsica* were collected from six populations chosen in order to cover the entire distribution range of the species. Because populations of this species are small, we collected leaf samples from all plants found in each population, for a total of 96 individuals (see Fig. 1 and Table 2 for details on each population). Leaf-tissue samples were dried and preserved in silica gel. Total genomic DNA was isolated from dried leaves using the QIAGEN^®^ DNeasy plant mini kit, following the manufacturer’s guidelines. Minor modifications were applied to the protocol and included an increased volume of buffer AP1 (from 400 μl to 600 μl), buffer AP2 (from 130 μl to 200 μl) and RNase A (from 4 μl to 6 μl), as well as a longer incubation time with buffer AP1 (30 minutes) for cell lysis.

### Microsatellite development and genotyping

DNA isolated from one specimen of *R. lamarmorae* sampled from population BC was used by Genetic Marker Services (Brighton, UK, www.geneticmarkerservices.com) to develop an enriched library, design and test microsatellite primer pairs. Enrichment involved incubating adaptor-ligated restricted DNA with filter-bonded synthetic repeat motifs, (AG)_17_, (AC)_17_, (AAC)_10_, (CCG)_10_, (CTG)_10_ and (AAT)_10_. Twenty-one positive library colonies were selected for sequencing, from which 15 microsatellites were designed and tested. The primer sets were designed using PRIMER 3.0 (Rozen & Skaletsky, 2000). Each primer pair was tested for amplification ability and polymorphism on eight individuals of *R. lamarmorae* (four from BC and four from SS). Cross-species amplification was tested on four individuals of *R. corsica* and four individuals of *R. chalepensis* (another species of the same genus with wider distribution also occurring in Sardinia; Salvo *et al*. 2010). Polymerase chain reactions (PCRs) for primer screening were performed in 25 μL and contained approximately 50 ng of DNA, 5 pmol of each unlabelled primer, 1.5 mM of MgCl_2_, 0.2 mM of each dNTP, 1X PCR buffer and 0.5 U of SupraTherm DNA Polymerase (GeneCraft, Cologne, Germany). *In-vitro* amplifications consisted of: 60 s denaturation at 95 °C, followed by 25 cycles of 95 °C for 60 s, annealing at 55 °C for 60 s and 72 °C for 60 s, ending with a final extension at 72 °C for 5 min. This first screening of microsatellites was performed on high-resolution agarose gels.

The primer pairs able to amplify polymorphic products were then used to genotype all sampled individuals via amplification with fluorescently labelled primers and separation of PCR products by capillary electrophoresis. Amplifications were performed following the two-step method described by Schuelke (2000), using ca. 20 ng of genomic DNA, 2.5 μl of 10X reaction buffer, 0.5 μl of each dNTP (10 mM), 1 μl of MgCl_2_ (50 mM), 0.2μl of the forward primer with M13(−21) tail at the 5’ end (10 μM), 0.5 μl of the reverse primer (10 μM), 0.5 μl of the fluorescently labelled M13(−21) primer (FAM, NED, VIC, PET; 10 μM) and 0.1 μl of Taq DNA polymerase (5 U/μl; Bioline GmbH, Luckenwalde, Germany) in a final volume of 25μl. Amplification reactions started with 94°C for 3 min, followed by 30 cycles of 94°C for 30 s,s 55°C for 45 s (see Table 1), and 72°C for 1 min. The fluorescently labelled M13(−21) primer was incorporated in the following eight cycles of 94°C for 30 s, 53°C for 45 s, and 72°C for 1 min, followed by a final extension step of 72°C for 5 min. Up to four PCR products of different primer sets were pooled for each individual and separated by capillary electrophoresis on an AB3130xl Genetic Analyzer. Alleles were sized against the internal size standard GeneScan^™^ LIZ500^™^ (Applied Biosystems) and scored using GeneMapper^®^ software Version 4.0 (Applied Biosystems). The observed allele size of all genotyped individuals was decreased by 18bp in order to account for the M13(−21) universal sequence tag.

**Table 1.**
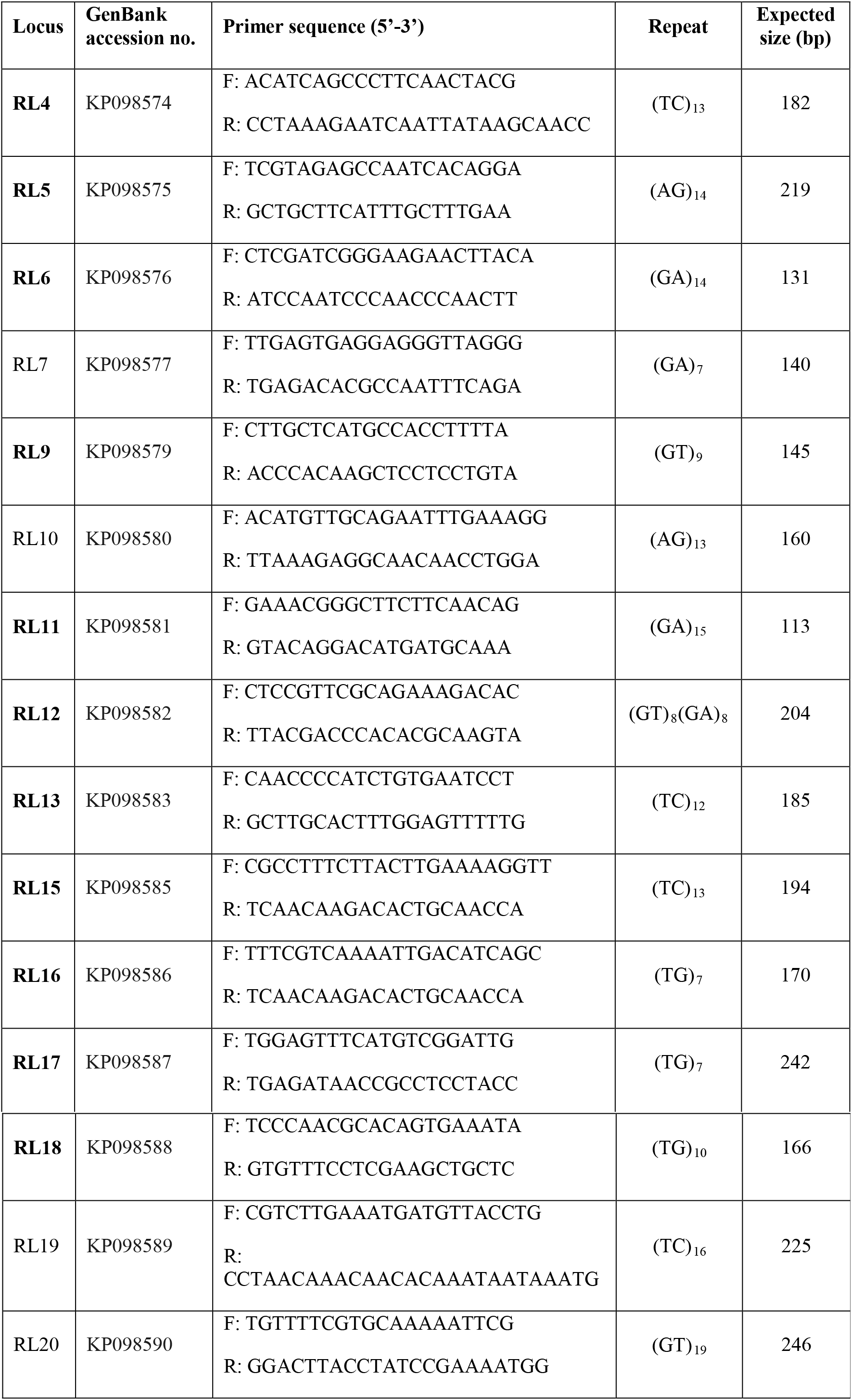
Characteristics of 15 microsatellite markers developed in *Ruta lamarmorae*. Shown for each marker are the GenBank accession number, forward and reverse sequences of the primer pair, repeat motif, and size of the expected PCR product (bp). Primers in bold were used for genotyping; locus RL9 was excluded from statistical analyses as all genotyped individuals of *R. corsica* and *R. lamarmorae* were heterozygotes for the same two alleles.

### Population genetic analyses

Even though *R. lamarmorae* is known to be tetraploid (Honsell 1957), a maximum of two alleles per locus and per individual were detected in all populations, thus showing disomic inheritance. Because genetic analyses can be performed with standard population genetic tools developed for diploid organisms in tetraploid species characterized by disomic inheritance (Stift *et al*. 2008; Meloni *et al*. 2013), our analyses were conducted assuming a diploid status for *R. lamarmorae*.

We assessed genetic diversity by quantifying the number of alleles (*N_A_*), proportion of polymorphic loci (*P*), number of private alleles (*A_P_*), observed (*H_o_*) and expected (*H_e_*) heterozygosity for each population across loci. Populations were tested for deviations from Hardy-Weinberg (HW) equilibrium using Fisher’s exact test with the Markov chain algorithm (Guo & Thompson 1992). The fixation index, *F*_IS_, was estimated in order to assess departure from Hardy-Weinberg expectations due to non-random mating. Fisher’s exact test was performed within each population to check for linkage disequilibrium (LD) between all different pairs of loci. These analyses were performed with the web-based Genepop (Raymond & Rousset 1995; Rousset 2008) and GenAlEx v.6.5 (Peakall & Smouse 2012). The program BOTTLENECK (Piry *et al*. 1999) was used to detect recent genetic bottlenecks in the studied populations. Based on the observed number of alleles and the sample size of a population, the program computes the gene diversity expected under the assumption of mutation-drift equilibrium and compares it to Hardy–Weinberg gene diversity (*H_e_*; Nei 1978) to establish whether there is a significant deficit of gene diversity resulting from a recent bottleneck. As recommended by the authors of the program, a Wilcoxon sign-rank test (Luikart *et al*. 1998) was performed using 1000 bottleneck simulation replicates under the stepwise mutation model (SMM).

FSTAT 2.9.3 (Goudet 1995) was used to estimate genetic differentiation among populations by *F*_ST_. We also measured *R*_ST_, an analogue of *F*_ST_ specific for microsatellite data, employing a stepwise mutation model (SMM, Slatkin 1995); *R*_ST_ was measured using the software SPAGeDi 1.4 (Hardy & Vekemans 2002). The same program was used to perform an allele-size permutation test (10000 permutations) to evaluate the contribution of stepwise mutations to the observed genetic differentiation, hence the relative suitability of *F*_ST_ vs. *R*_ST_ (Hardy *et al*. 2003). In addition, since *F*_ST_ can considerably underestimate differentiation when loci are highly variable (as commonly found with microsatellite markers; Hedrick 2005; Jost 2008; Meirmans & Hedrick 2011), we also calculated Jost’s *D*_est_ (Jost 2008) using 1,000 bootstrap replicates in SMOGD (Crawford 2010).

An Analysis of Molecular Variance (AMOVA) was performed to assess the hierarchical partitioning of genetic variation among populations and species. We followed the procedure of Excoffier *et al*. (1992), Huff *et al*. (1993), Peakall *et al*. (1995), and Michalakis & Excoffier (1996) by estimating *F*_ST_ and using 999 random permutations of the data in GenAlEx v.6.5 (Peakall & Smouse 2012). With the same program, a Principal Coordinate Analysis (PCoA) based on a genetic distance matrix was performed to visualise genetic relatedness among individuals. To test for isolation by distance, a Mantel test (Mantel 1967) was applied to the matrix of pairwise Nei’s genetic distances (Nei 1972, 1978) and the matrix of geographical distances with 999 random permutations in GenAlEx v. 6.5 (Peakall & Smouse 2012).

Population structure was inferred using the Bayesian clustering method implemented in STRUCTURE (v. 2.3; Pritchard *et al*. 2000; Hubisz *et al*. 2009). The program uses a Markov Chain Monte Carlo (MCMC) procedure to estimate P(*X|K*), the posterior probability that the data fit the hypothesis of *K* clusters, and assigns individual genotypes to clusters by estimating the membership coefficient Q for each individual based on allele frequencies at unlinked loci (independent of locality information). We tested all possible values of *K* from 1 to 9; for each *K* we ran an admixture model (each individual draws some fraction of the genome from each of the *K* populations) with correlated allele frequencies 20 times with a length of burnin period of 100,000 followed by 100,000 MCMC repetitions. To identify the best *K*, we measured ΔK (the rate of change in the log probability of data between successive *K* values), as suggested by Evanno *et al*. (2005) and implemented in STRUCTURE HARVESTER (Earl & vonHoldt 2012). This method provides the most accurate estimate of the number of clusters *K* (Evanno *et al*. 2005), but does not allow for discrimination between K=1 and K=2. Therefore we also calculated the average posterior probability of the data for each value of *K*, Ln P(*X|K*), as proposed by Pritchard *et al*. (2000). After determining the most effective number of genetic groups (*K*) for our data, we ran STRUCTURE with the admixture model and default parameter settings; the inferred genetic composition of individuals was then determined using 100,000 iterations after a burnin period length of 100,000.

## Results

### Microsatellite development and cross-species amplification

Of the 15 microsatellites newly developed for *R. lamarmorae*, 13 were suitable for genetic analyses on the target species (Online Resource 2). Tests on cross-species amplification showed that 13 markers could be used for *R. corsica* while 11 could be amplified in the more distantly related *R. chalepensis* (Online Resource 2).

### Genetic diversity

Eleven microsatellites showing a clear amplification pattern after capillary electrophoresis on both *R. corsica* and *R. lamarmorae* were used for genotyping (Table 1). Because all studied individuals of *R. corsica* and *R. lamarmorae* were heterozygotes for the same two alleles in locus RL9, this marker was excluded from population genetic analyses. In *R. lamarmorae*, all amplified loci were polymorphic; in *R. corsica*, locus RL16 was fixed in populations SA, GH, AL, BA; population SA showed fixed alleles also at loci RL15, RL17 and RL18. The number of alleles identified across all loci ranged from 22 to 61 in *R. corsica* populations, and from 55 to 68 in the two *R. lamarmorae* sub-populations (Table 3). Private alleles were found in all populations except SA (Table 2).

**Table 2.** Description of populations of *R. corsica* and sub-populations of *R. lamarmorae* surveyed in this study.

| Species | Population abbreviation | Location | Population size | Sample number | Coordinates | Altitude |
| --- | --- | --- | --- | --- | --- | --- |
| *R. lamarmorae* | BC | Broncu Spina | ~1000 | 30 | 40° 07' 34" N<br>9° 31' 26" E | 1650m |
|  | SS | Su Susciu | ~1000 | 30 | 40° 01' 02" N<br>9° 19' 33" E | 1620m |
| *R. corsica* | MU | Muvrella | 20 | 8 | 42° 24' 0" N<br>8° 54' 0" E | 1050m |
|  | MC | Monte Cinto | 150 | 16 | 42° 23' 0" N<br>8° 55' 0" E | 1750m |
|  | SA | Saltare | 20 | 8 | 42° 21' 41" N<br>8° 52' 53" E | 1180m |
|  | GH | Ghisoni | 40 | 23 | 42° 6' 37" N<br>9° 9' 16" E | 1650m |
|  | AL | Albertacce | 120 | 20 | 42° 17' 39" N<br>8° 52' 44" E | 1300-1500m |
|  | BA | Bastelica | 750 | 21 | 42° 0' 26" N<br>9° 6' 4" E | 900m |

**Table 3.** Genetic diversity parameters inferred from microsatellite analysis. *N_A_* total number of alleles across loci, *P* proportion of polymorphic loci, *A_P_* number of private alleles, *H_o_* observed heterozygosity, *H_e_* expected heterozygosity, *F*_IS_ fixation index; SD, standard deviation. For abbreviations of populations see Table 1.

| Population | $N_A$ | $P$ | $A_P$ | $H_o \pm SD$ | $H_e \pm SD$ | $F_{IS} \pm SD$ |
| --- | --- | --- | --- | --- | --- | --- |
| BC | 55 | 1 | 4 | $0.554 \pm 0.082$ | $0.614 \pm 0.078$ | $0.097 \pm 0.100$ |
| SS | 68 | 1 | 8 | $0.587 \pm 0.076$ | $0.639 \pm 0.083$ | $0.082 \pm 0.027$ |
| Overall *R. lamarmorae* | 123 | 0.967 | 12 | $0.571 \pm 0.055$ | $0.627 \pm 0.056$ | $0.109 \pm 0.052$ |
| MU | 51 | 1 | 2 | $0.623 \pm 0.075$ | $0.688 \pm 0.072$ | $0.094 \pm 0.056$ |
| MC | 55 | 1 | 3 | $0.578 \pm 0.077$ | $0.630 \pm 0.073$ | $0.081 \pm 0.076$ |
| SA | 22 | 0.6 | 0 | $0.350 \pm 0.118$ | $0.323 \pm 0.105$ | $-0.083 \pm 0.073$ |
| GH | 53 | 0.9 | 1 | $0.546 \pm 0.082$ | $0.593 \pm 0.084$ | $0.080 \pm 0.041$ |
| AL | 61 | 0.9 | 1 | $0.624 \pm 0.085$ | $0.624 \pm 0.086$ | $-0.001 \pm 0.031$ |
| BA | 53 | 0.9 | 3 | $0.497 \pm 0.080$ | $0.533 \pm 0.081$ | $0.070 \pm 0.061$ |
| Overall *R. corsica* | 295 | 0.883 | 10 | $0.536 \pm 0.036$ | $0.543 \pm 0.035$ | $0.160 \pm 0.024$ |

All populations, except for GH, were at HW equilibrium (p>0.05). *F*_IS_ values were positive in the two sub-populations of *R. lamarmorae*, as well as in four out of six populations of *R. corsica* (MU, MC, GH and BA; Table 3), meaning that the departure of genotype frequencies from Hardy-Weinberg expectations was always associated with a deficit of heterozygotes. Conversely, an excess of heterozygotes was found in populations SA and AL (*R. corsica;* Table 3).

Significant linkage-disequilibrium (LD) at the 5% level was detected at one pair of loci for populations GH (RL6-RL18), AL (RL13-RL15) and MU (RL12-RL17), three loci for population BC (RL4-RL6, RL13-RL15, RL11-RL17), four loci for SA (RL4-RL12, RL4-RL13, RL11-RL13, RL12-RL13) and BA (RL6-RL11, RL12-RL15, RL13-RL15, RL15-RL18), and eight loci for SS (RL4-RL5, RL5-RL11, RL5-RL13, RL6-RL13, RL11-RL13, RL15-RL17, RL12-RL18, RL15-RL18). It was impossible to perform most of the tests for populations MU (25 out of 45 pairs of loci) and SA (30 out of 45 pairs of loci).

Gene diversity (*H_e_*) was high in all populations, with a mean value of 0.627 for *R. lamarmorae* and 0.543 for *R. corsica;* the only population showing a relatively low value was SA (*R. corsica*), with *H_e_*= 0.323 (Table 3).

The only population showing significant signs of a recent bottleneck was BA (*R. corsica*, p = 0.032), while all other populations were at mutation-drift equilibrium.

### Genetic structure

Genetic differentiation among populations measured with *F*_ST_ was always statistically significant (*P* < 0.05); it was 0.086 between the two sub-populations of *R. lamarmorae* and ranged between 0.012 (MU-MC) and 0.240 (AL-SA) in *R. corsica* (Table 4). Genetic differentiation among populations across the two species ranged between 0.050 (MU-SS) and 0.212 (SA-BC). Population SA was the most differentiated, with 0.173<*F*_ST_<0.240 (Table 4). The overall genetic differentiation among populations was significant (p=0.01), with *F*_ST_=0.097 for *R. corsica* and *F*_ST_=0.086 for *R. lamarmorae*. Similarly, the AMOVA (based on *F*_ST_) displayed that most of genetic variation was partitioned within-population (88%), while only 9% was found among populations and 3% between islands (Fig. 2). Values of *R*_ST_ (analogous to *F*_ST_, but specific to microsatellite data; Slatkin 1995) were slightly smaller than those of *F*_ST_ (0.003< *R*_ST_ <0.253) (Online Resource 3). However, observed *R*_ST_ values were not significantly higher than permuted *R*_ST_, suggesting that stepwise mutations did not contribute to population differentiation and that *F*_ST_ (based on allele identity) explains genetic differentiation among populations better than *R*_ST_ (based on allele-size information; Hardy *et al*. 2003). Although *F*_ST_ is widely used as a measure of population structure, its dependency on expected heterozygosity might underestimate the genetic differentiation among populations. Higher levels of genetic differentiation among populations were detected, in fact, when the partition of diversity was based on the effective number of alleles (i.e. *D*_est_, Jost 2008) rather than on the expected heterozygosity. We found *D*_est_=0.179 between the two sub-populations of *R. lamarmorae*, and 0.003 <*D*_est_<0.238 for *R. corsica* (Table 4). Across populations, total *D*_est_ for *R. corsica* was 0.162.

**Fig. 2.**
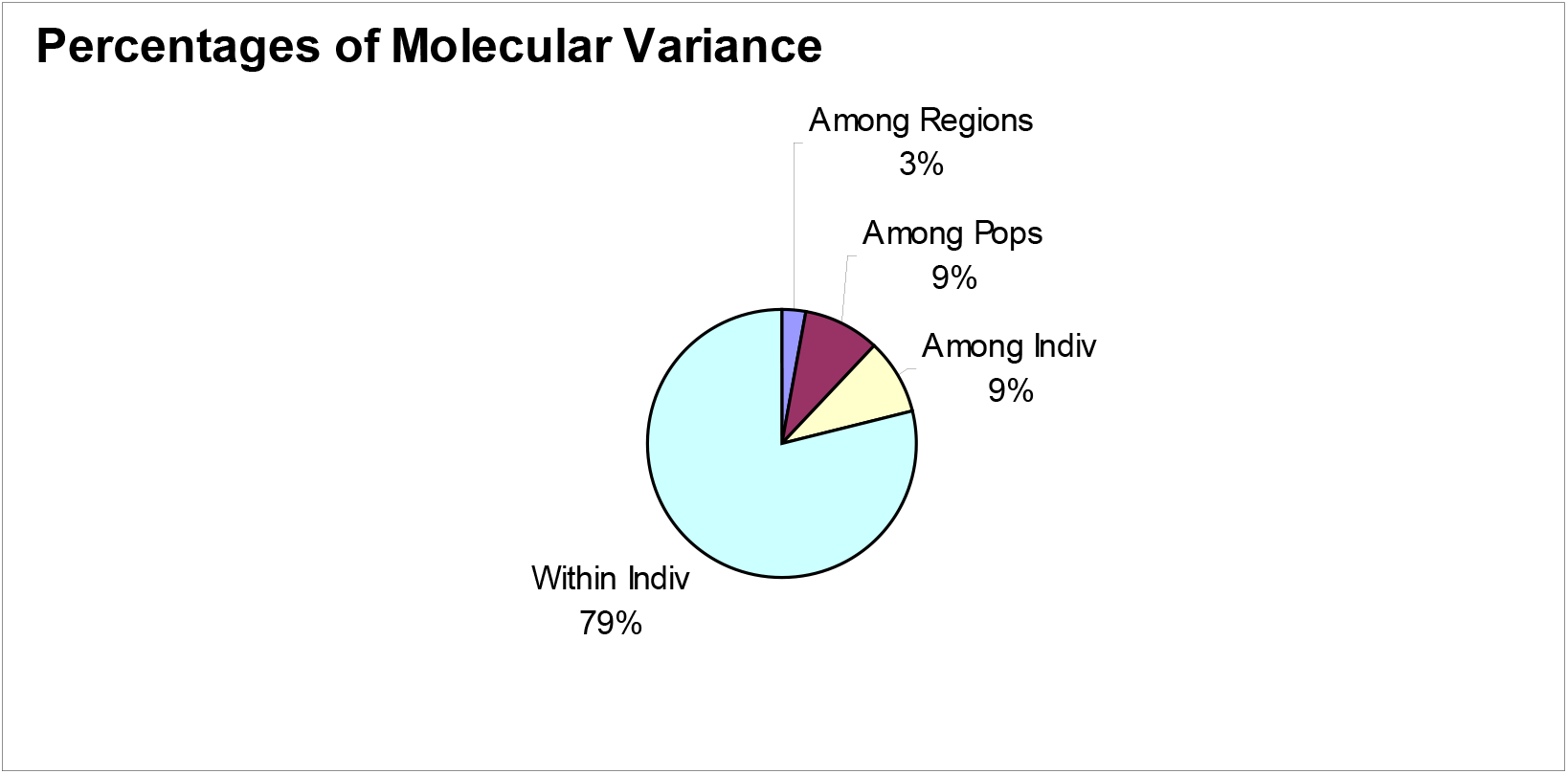
Analysis of Molecular Variance (AMOVA) based on ten microsatellite markers used to genotype 96 individuals taxonomically assigned to *R. corsica* (six populations) and 60 individuals taxonomically assigned to *R. lamarmorae* (one population divided in two sub-populations). The two regions considered in this analysis represent the two islands.

**Table 4.** Pairwise population estimates of *F*_ST_ (diagonal below) and *D*_est_ (above diagonal). All values are statistically significant (95% CI). For abbreviations of populations see Table1.

|  | *R. lamarmorae* |  | *R. corsica* |  |  |  |  |  |
| --- | --- | --- | --- | --- | --- | --- | --- | --- |
|  | BC | SS | MU | MC | SA | GH | AL | BA |
| BC |  | 0.179 | 0.146 | 0.155 | 0.303 | 0.234 | 0.256 | 0.254 |
| SS | 0.086 |  | 0.278 | 0.121 | 0.208 | 0.156 | 0.099 | 0.193 |
| MU | 0.087 | 0.050 |  | 0.003 | 0.177 | 0.076 | 0.065 | 0.103 |
| MC | 0.102 | 0.065 | 0.012 |  | 0.179 | 0.048 | 0.060 | 0.077 |
| SA | 0.212 | 0.173 | 0.176 | 0.173 |  | 0.168 | 0.238 | 0.158 |
| GH | 0.130 | 0.083 | 0.052 | 0.038 | 0.184 |  | 0.131 | 0.124 |
| AL | 0.129 | 0.060 | 0.046 | 0.044 | 0.240 | 0.085 |  | 0.156 |
| BA | 0.160 | 0.136 | 0.092 | 0.071 | 0.213 | 0.088 | 0.122 |  |

The PCoA showed some genetic structure among *R. corsica* populations: while MU, MC and GH appeared strongly genetically interconnected, AL and BA overlapped only slightly, and SA was well separated from all other populations (Fig. 3A). A clear genetic differentiation was also present between the two sub-populations of *R. lamarmorae* (Fig. 3B). When considering both species together, population structure was still clearly present, although less marked: the two species were well separated, populations within each species overlapped slightly more, and population SA, taxonomically assigned to *R. corsica*, remained well separated from all remaining populations (Fig. 3C).

The Mantel test for correlation between genetic differentiation and geographic distances among populations was not significant (p=0.579, R^2^ =0.0076).

**Fig. 3.**
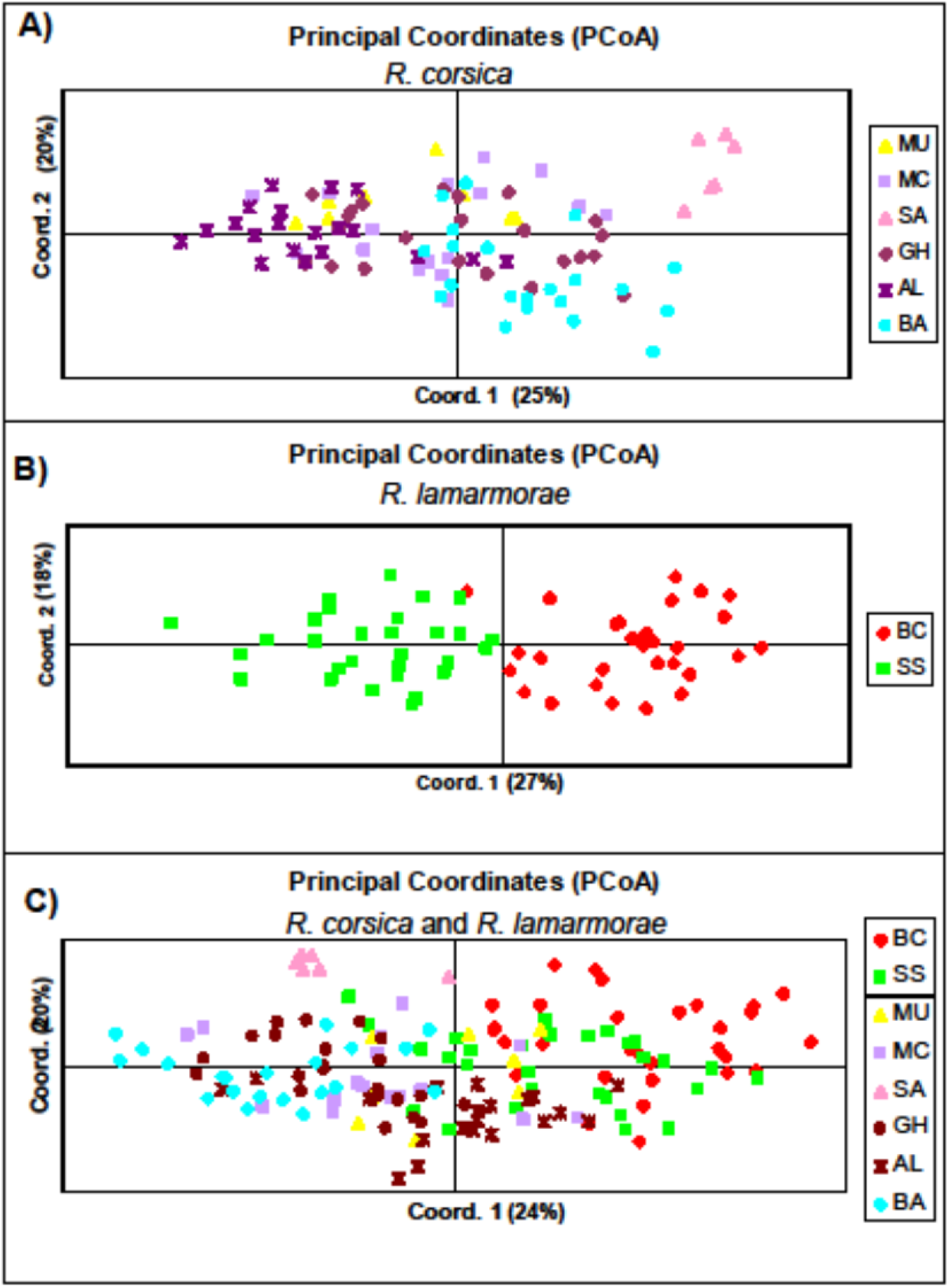
Principal Coordinate Analysis (PCoA) based on the multilocus genotypes of 96 individuals of *R. corsica* (A), 60 individuals of *R. lamarmorae* (B), and the two species together (C). The percentage of the total variability explained by the first two components is indicated on brackets. Each symbol represents a single plant from one of the eight studied populations. Information on each population is provided in Table 2.

For both *R. corsica* and *R. lamarmorae* the Ln probability of the data (Ln P(*X|K*)) did not allow for discrimination between different values of *K* (Table S3) while, based on the Δ*K* statistic, the best-supported number of *a posteriori* genetic clusters was *K*=3 for *R. corsica* (Online Resource 1 A) and *K*=2 for *R. lamarmorae* (Online Resource 1 B). For *K*=3, populations of *R. corsica* MU, MC, GH and AL were grouped together, while SA and BA appeared genetically distinct (Fig. 4A). Very little admixture seemed to occur between the two *R. lamarmorae* sub-populations for *K*=2 (Fig. 4B). When *R. corsica* and *R. lamarmorae* were pooled together, the most effective number of genetic groups was *K*=4 according to Pritchard’s method (Pritchard *et al*. 2000, Table S3) and *K*=2 according to Evanno’s method (Evanno *et al*. 2005, Online Resource 1 C). Figure 5 shows subfigures for *K*=4 and *K*=2; subfigures were included also for *K*=3 and 5<*K*<7, because populations subdivision was informative also for these values of *K*. For *K*=2, populations of the two species were well separated in two distinct *R. corsica* and *R. lamarmorae* clusters, except for population SA, taxonomically classified as *R. corsica*, whose individuals were genetically assigned with higher probability to populations of *R. lamarmorae* (Fig. 5). Values of *K*=3 showed the separation of the two *R. lamarmorae* sub-populations, while population SA (of *R. corsica)* clustered with sub-population SS (of *R. lamarmorae;* Fig. 5). For *K*=4, *R. lamarmorae* sub-populations were genetically distinct, population SA was separated from all populations, while the remaining populations of *R. corsica* formed a single cluster (Fig. 5). Values of *K*=5 showed the separation of the *R. corsica* population BA (Fig. 5). Values of *K*=6 and *K*=7 showed more admixture. It is notable that population SA, taxonomically assigned to *R. corsica*, remained separated from all other populations for all *K* values (Fig. 5)

**Fig. 4.**
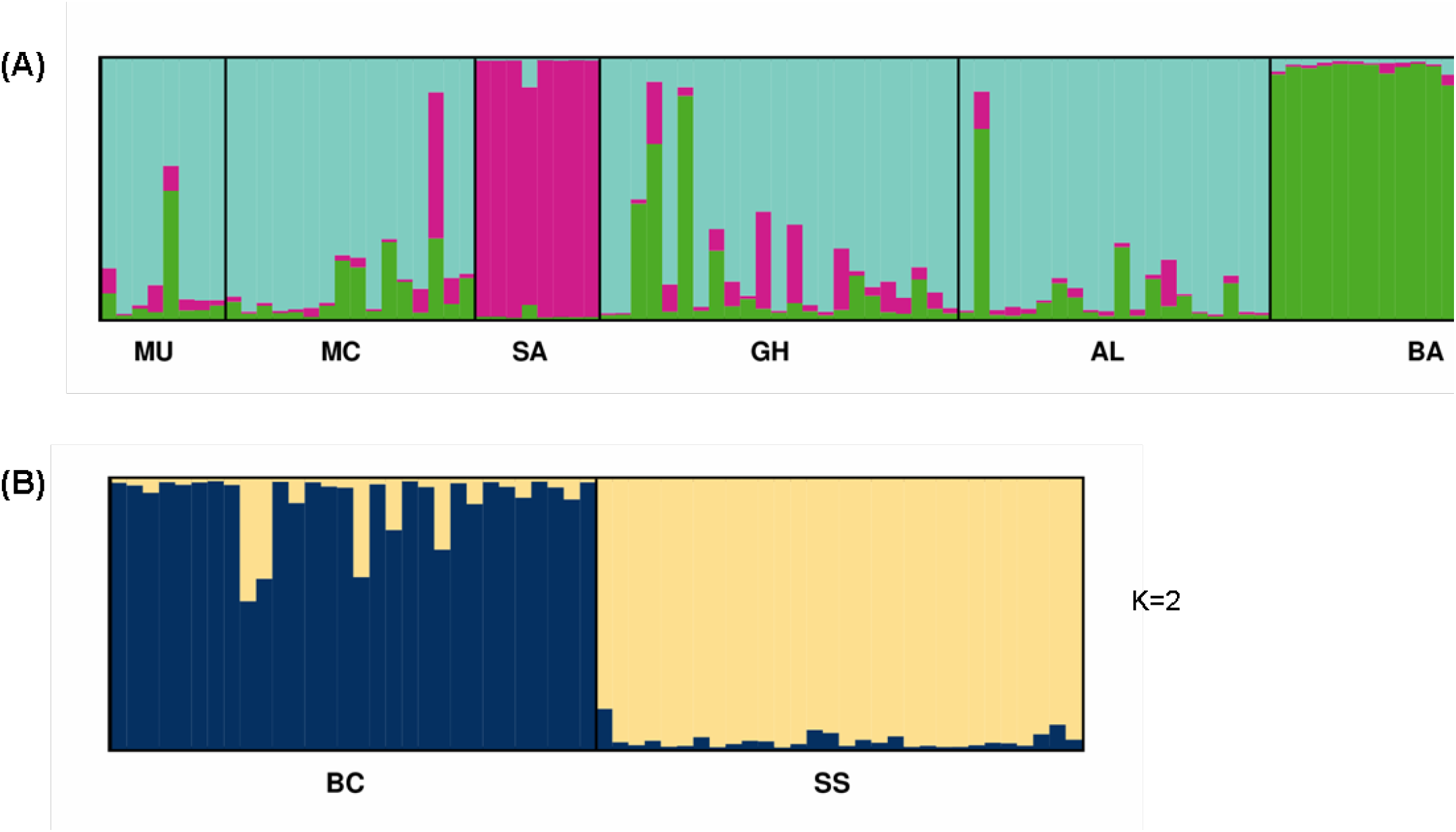
Proportional membership (Q) to clusters identified by STRUCTURE for: (A) 96 individuals from the six studied populations *of R. corsica;* (B) 60 individuals from the two studied sub-populations of *R. lamarmorae*. Colours distinguish between the different clusters identified for each of the above three scenarios. Each individual is represented by a single vertical bar and grouped by locality, indicated on the x-axis. Information on each population (locality) is provided in Table 2.

**Fig. 5.**
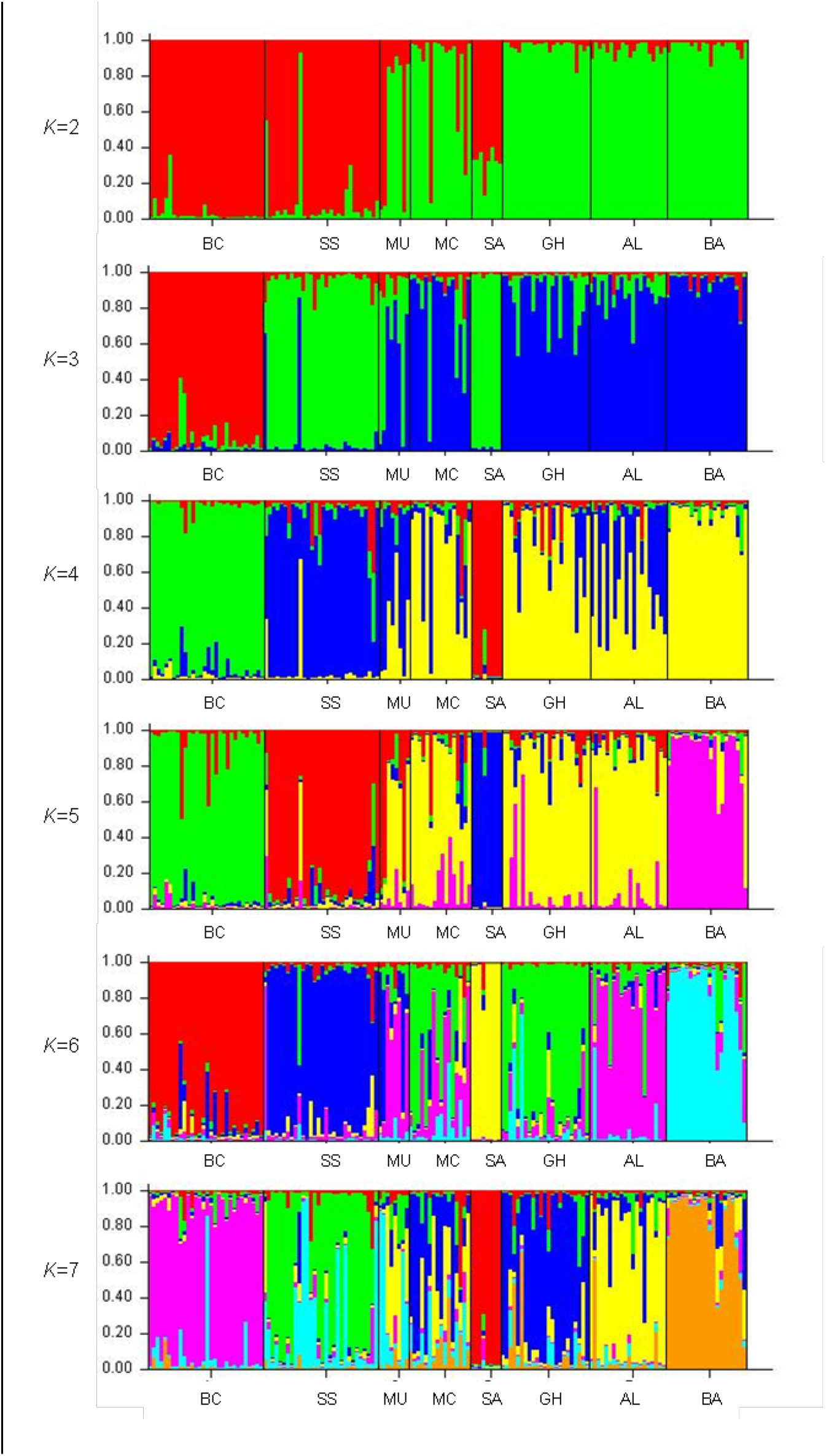
Proportional membership (*Q*) to clusters identified by STRUCTURE for 2<*K*<7. Colours distinguish between the different *K* clusters. All studied individuals of *R. lamarmorae* (sub-populations BC and SS: 60 individuals) and *R. corsica* (populations MU, MC, SA, GH, AL and BA: 96 individuals) are represented by vertical bars and grouped by locality, indicated on the x-axis. Information on each population (locality) is provided in Table 2.

## Discussion

### Microsatellite development and cross-species amplification

Most of the microsatellites developed for this study were suitable for genetic analyses on *R. lamarmorae* (13 out of 15), *R. corsica* (13 out of 15) and the more distantly related *R. chalepensis* (11 out of 15), being amplifiable and polymorphic (Online Resource 2). Despite the difference in ploidy level between *R. corsica* (diploid) and *R. lamarmorae* (tetraploid), the high transferability of the molecular markers suggests a low divergence between the genomes of the two species.

### Genetic diversity

Among the microsatellite markers developed for the present study, 11 were useful for population genetic analyses on *R. corsica* and *R. lamarmorae*. Our study revealed high levels of genetic diversity in both *R. corsica* (*P*=0.883, *H_e_*=0.543) and *R. lamarmorae* (*P*=0.972, *H_e_*=0.627). Even though each species is restricted to a single island (Fig. 1), most populations showed fewer genetic signatures of isolation and small population size than typically expected for insular endemics (i.e., low gene diversity, high homozygosity, high LD; Ellstrand 1993, Frankham 1998, Bouzat 2010). Unexpectedly high values of genetic diversity were also found in the congener *R. oreojasme*, an endangered species endemic to the island of Gran Canaria in the Canarian archipelago (*A*=7.625, *P*=0.984, *H_o_*=0.558, *H_e_*=0.687; Meloni *et al*. 2015).

The occurrence of another endangered, insular endemic with high genetic diversity within the same genus suggests that *R. corsica* and *R. lamarmorae* share some intrinsic traits maintaining high levels of genetic variation with *R. oreojasme*. Hermaphroditism and proterandry, characterising all three endemics, favour outcrossing, thus increasing genetic diversity. Furthermore, polyploidy, occurring in all *Ruta* species except *R. corsica*, might also contribute to elevated genetic variation, for polyploids are usually characterised by higher genetic diversity than diploids (Soltis & Soltis 2000). Hermaphroditism, proterandry and polyploidy were associated with high levels of genetic variation also in other insular endemics outside of *Ruta* (Pérez de Paz & Caujapé-Castells 2013). Conversely, genetic analyses of *R. microcarpa*, a critically endangered species endemic to the island of La Gomera (also in the Canarian archipelago), revealed levels of genetic diversity lower than those observed in *R. corsica, R. lamarmorae* and *R. oreojasme* (*H_o_*=0.651, *H_e_*=0.410; Meloni *et al*. 2014). The relatively low genetic diversity found in *R. microcarpa*, resulting mainly from the almost total absence of sexual reproduction in this species (Meloni *et al*. 2015), highlights the role of outcrossing in maintaining variation within and among populations.

Despite the relatively low number of population genetic studies on C-S species, a general trend of high genetic diversity was detected in published analyses of Corsican and/or Sardinian endemics. High levels of genetic variability were found, for example, in the Sardinian *Centaurea horrida* (*H_e_*=0.780, Mameli *et al*. 2008) and *Centaurea filiformis* (*H_e_*=0.678, Pisanu *et al*. 2011; *H_e_* = 0.576; Giorgio Binelli, pers. comm.), in the Corso-Sardinian *Ferula arrigoni* (*P*=0.92, *H_w_* = 0.317, Dettori *et al*. 2014b), and in the Corsican *Mercurialis corsica* (Migliore *et al*. 2011).

The occurrence of high genetic diversity in several C-S species suggests that also some island-dependent factors (i.e., age, physical dimension, geologic origin and history of the islands) might have contributed to the genetic diversity of *R. corsica* and *R. lamarmorae* as well as of several other C-S endemics.

The complex geologic history of Corsica and Sardinia most likely influenced the evolution and current genetic diversity of their endemics in different ways. Corsica and Sardinia are considered old islands (they broke off from the Iberian peninsula as the C-S microplate ca. 30-28 MYA) and have been proposed to harbour many relictual endemics (Médail & Quézel 1999, Grill *et al*. 2007). Because they are continental fragment islands, it has been suggested that they might have hosted the ancestors of many of their current endemic taxa before their separation from the Iberian peninsula in the Oligocene, although this hypothesis has been corroborated by molecular dating analyses only in a few cases (for example, in Araceae; Mansion et al., 2008). Subsequent to its split from the Iberian peninsula, the C-S microplate was temporarily connected with the Apulian microplate during the Miocene. The vicariant origin of some C-S species during the Oligocene and Miocene might have favoured relatively high levels of genetic diversity, because they did not experience the loss of genetic variation associated to LDD and founder events typical, for example, of oceanic island colonization. Furthermore, the long time-spans since colonisations (from either the Iberian peninsula in the Oligocene or the Apulian microplate in the Miocene) might have increased opportunities to accumulate genetic variation through mutation, recombination, drift and selection in CS endemics. Additionally, multiple overland migrations might have occurred while Corsica and Sardinia were connected to neighbouring continental masses via land bridges at different points in time during the Miocene and Pleistocene, thus favouring genetic exchanges also within species with limited dispersal ability, as in the case of *R. corsica* and *R. lamarmorae* (Gianluigi Bacchetta, pers. obs.).

Furthermore, the relatively stable climate characterising Corsica and Sardinia during Quaternary glacial/interglacial periods (Taberlet 1998, Grill *et al*. 2007) might also have contributed to the high genetic diversity detected in their endemics, for it allowed populations of many species to persist through several climatic cycles and accumulate genetic differences via mutation and recombination. Finally, climatic changes during the Miocene and Pleistocene might have also contributed to the high genetic variation detected in C-S species. Indeed, land bridges among Corsica, Sardinia, the Apulian plate and the African continent during the MSC (Hsü *et al*. 1977, Krijgsman *et al*. 1999, McKenzie 1999) and Quaternary glacial marine regressions (Thompson 2005) may have decreased the genetic effects of isolation by favouring genetic exchange between C-S populations and populations of the same species occurring on other land masses. Among the above mentioned island-dependent factors, the vicariant origin of the ancestor of *R. corsica* and *R. lamarmorae* following the MSC, supported by molecular dating and biogeographic analyses (Salvo et al., 2010), and the climatic stability of Corsica and Sardinia during Pleistocene glacial cycles might have contributed to the relatively high levels of genetic diversity detected in the two species.

*Ruta lamarmorae* showed slightly higher values for most diversity parameters than *R. corsica* (*P*=0.967, *H_o_*=0.571, *H_e_*=0.627 vs. *P*=0.83, *H_o_*=0.536, *H_e_*=0.543). As mentioned above, the higher ploidy of the former might explain this difference. In addition to intrinsic traits, the small size of most *R. corsica* populations (Laetitia Hugot, pers. obs.) and its scattered distribution could also explain the lower diversity found in *R. corsica*, because its populations might be more affected by inbreeding and isolation than the larger population of *R. lamarmorae* (Hamrick & Godt 1989; Ouborg *et al*. 2006).

An exception to the trend of relatively high genetic diversity found in *R. corsica* and *R. lamarmorae* is represented by population SA (taxonomically assigned to *R. corsica*), which showed some genetic erosion, being characterized by values of polymorphism, allelic diversity and heterozygosity much lower than the other populations analysed in this study (*N_A_* =22, *P*=0.6, *H_o_*=0.350, *H_e_*=0.323, Table 3). The low genetic diversity and the presence of alleles fixed at four loci suggest that genetic drift and inbreeding strongly influenced the genetic structure of this small population (20 individuals in total; Laetitia Hugot, unpublished data; Ellstarnd *et al* 1993). They also suggest that the rate of genetic exchange between SA and other *R. corsica* populations is extremely low, and it seems unable to counteract the genetic effects of small population size.

### Population genetic structure

Genetic differentiation among populations measured with *F*_ST_ and AMOVA (Table 4 and Fig. 2 respectively) was lower than expected, when considering the low dispersal ability of *R. corsica* and *R. lamarmorae* (Gianluigi Bacchetta, unpublished data). Because *F*_ST_ is strongly influenced by levels of heterozygosity (Meirmans & Hedrick 2011), we cannot rule out the possibility that *F*_ST_ values (and AMOVA, accordingly) may simply reflect the high levels of heterozygosity detected in our study species (Table 3), and that in this case *D*_est_ (which is based on the effective number of alleles and thus independent of levels of heterozygosity; Jost 2008; Table 4) might better describe genetic differentiation among populations. Indeed, *D*_est_ values are approximately two times larger than *F*_ST_ values (*F*_ST_=0.097, *D*_est_=0.162 for *R. corsica; F*_ST_=0.086, *D*_est_=0.179 for *R. lamarmorae*), suggesting that the studied populations might be more differentiated than implied by *F*_ST_. High values of heterozygosity and lower levels of *F*_ST_ than *D*_est_ were also found in the congener *R. oreojasme* (*H_e_*= 0.687, *F*_ST_ = 0.097, *D*_est_ = 0.275; Meloni *et al* 2015).

The differentiation among populations described by *D*_est_ (high for *R. lamarmorae*, relatively lower for *R. corsica;* Table 4) was supported by PCoA (Fig. 3) and Bayesian analyses (Fig. 4 and 5). The population structure detected in the two species (Fig. 3, 4 and 5) and the absence of isolation by distance reiterates that limited gene flow probably due to low dispersal ability has played a significant role in shaping current patterns of genetic differentiation. Low dispersal ability seems to characterise particularly *R. lamarmorae*, whose sub-populations, showing the highest value of within-species *D*_est_ and appearing genetically well separated in PCoA and Bayesian analyses (Fig. 3B and 4), are only ca. 3 km apart from each other, Analyses that included both species indicated some genetic differentiation between *R. corsica* and *R lamarmorae* (Table 4, Fig 3C and 5), lending some support to their taxonomic classification as separated species. However, some degree of admixture between them was detected. The difference in ploidy level (*R. corsica* is diploid, *R. lamarmorae* is tetraploid; Contandriopoulos 1957, Honsell 1957), the low dispersal ability of the two species (Gianluigi Bacchetta, unpublished data), and their geographic separation through the Bonifacio strait suggest that relatively low levels of admixture detected might not be the effect of gene flow between islands. Instead, this result might be better explained by the possibility that *R. corsica* and *R. lamarmorae* diverged at only few loci, probably as a consequence of low mutation rate and/or the long generation time that characterises the two species. Further analyses would be required to test this hypothesis. The high transferability of the newly developed microsatellite markers between the two species (Online Resource 2) supports the hypothesis of low divergence.

In all genetic analyses, population SA (of *R. corsica*) strongly differed from all other studied populations (Table 4), forming a distinct group in both PCoA (Fig 3A and 3C) and STRUCTURE analyses (Fig 4 and 5). This result reinforces the conclusion that SA is more genetically isolated than the other studied populations. Because geographic distance does not explain this genetic isolation, other factors seem to affect gene flow in this population. Interestingly, in STRUCTURE analyses that included *R. corsica* and *R. lamarmorae*, population SA showed evidence of genetic admixture between the two species for *K*=2 (the most accurate estimate of the number of clusters obtained measuring Δ*K*; Evanno *et al*. 2005), was assigned to the same genetic group of *R. lamarmorae* sub-population SS for *K*=3, and resulted genetically differentiated from all other studied populations of both species for 4<*K*<7 (Fig. 5). Further genetic and karyotype studies are required to clarify the reasons of the genetic differentiation of SA. Genetic analyses on the congener *R. chalepensis*, occurring in both islands, might also help to clarify the occurrence and amount of gene flow between Corsica and Sardinia.

### Conservation implications

This study provides important insights into the genetic structure of *R. corsica* and *R. lamarmorae*, with potential applications for their effective conservation. The medium to high genetic diversity characterising these endemics suggests that both species are not at high risk of extinction due to genetic factors. Nevertheless, their isolation, their very restricted distribution, the small size of *R. corsica* populations and, in the case of *R. lamarmorae*, the continued anthropogenic and environmental threats to its population (i.e., overgrazing, fires, presence of skiing infrastructures; Bacchetta *et al*. 2006) might jeopardize conservation. Moreover, mountain habitats are considered particularly sensitive to climatic change and are likely to show the effects of temperature increase earlier and more markedly than other ecosystems (Grabherr *et al*. 2000; Thuiller *et al*. 2005; Patsiou *et al*. 2014). Germination tests in *R. lamarmorae* show that an increase in winter temperatures could lead to reduced reproductive capability, because seeds may not experience the vernalization period required for germination (Bacchetta *et al*., unpublished data). We expect a similar negative effect on germination in *R. corsica*, given the close relatedness with *R. lamarmorae*, the phenotypic traits they share, and the shared ecological preference for mountain habitats. Similarly, a reduction in seed germination rate with increasing temperatures has been observed in other Sardinian taxa that occur at medium to high altitudes, including *Lamyropsis microcephala* (Mattana *et al*. 2009), *Ribes multiflorum* subsp. *sandalioticum* (Mattana *et al*. 2012), and *Vitis vinifera* subsp. *sylvestris* (Orrù *et al*. 2012).

*In situ* conservation is essential to the long-term persistence of these two island endemics and should be aimed at preserving all extant populations, for they all carry unique alleles (a particular case is represented by population SA, whose low genetic diversity and high differentiation require further genetic analyses; Table 3 and 4, Fig. 3 and 5). For *R. lamarmorae*, seeds from sub-population BC were already collected for long-term storage at the BG-SAR (the Sardinian Germplasm Bank, University of Cagliari). Given the high differentiation detected between sub-populations BC and SS of *R. lamarmorae*, germplasm collection should be planned also from SS. Genetic studies on the eastern part of the Gennargentu massif are also advisable, because sub-populations from this area might show genetic differentiation as high as that found in the two *R. lamarmorae* sub-populations analysed here. Finally, more detailed studies on the reproductive biology and dispersal ability of these species are fundamental for planning specific, and thus potentially successful, conservation programs and guarantee their long-term survival.

---

**Online Resource 1.**
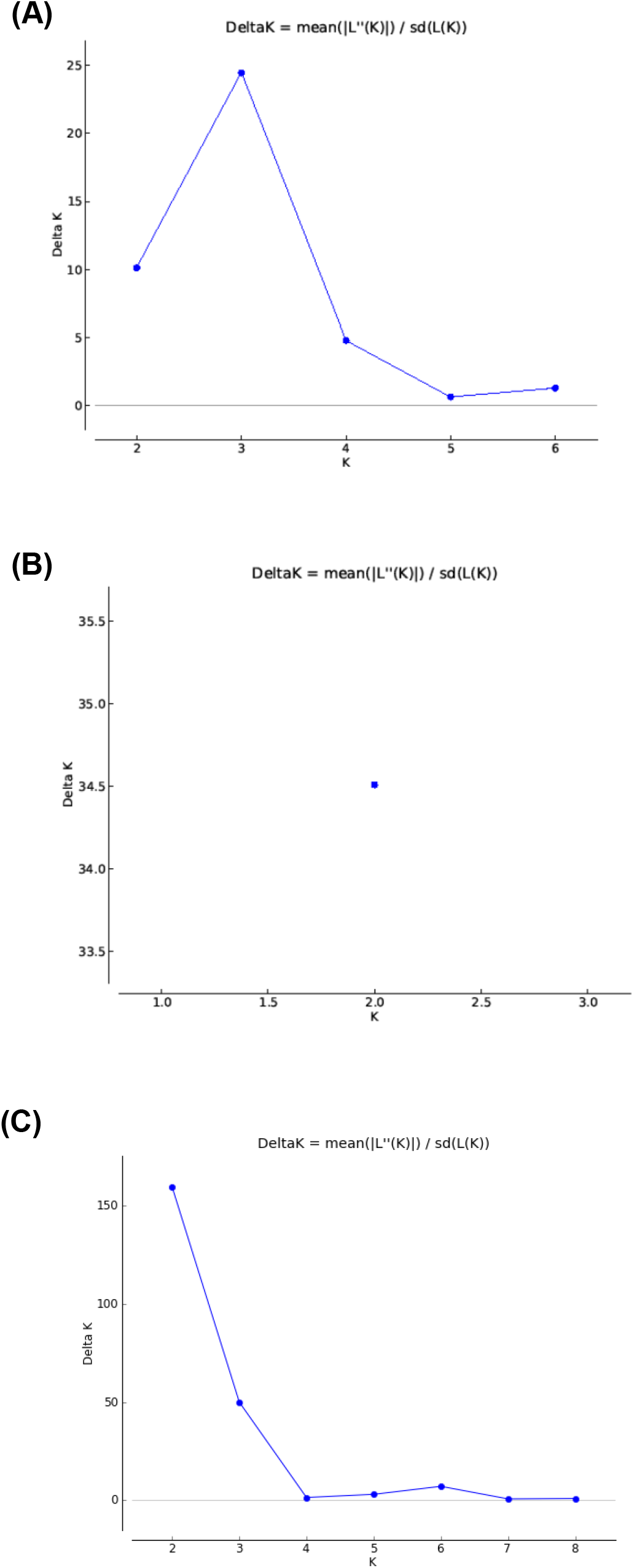
Δ*K* values (a measure of the rate of change in the structure likelihood function) as a function of *K* (the number of putative populations) for: (A) the six studied populations *of R. corsica*; (B) the two studied sub-populations *of R. lamarmorae* and (C) the two species analysed together. The model run here is an admixture model with correlated allele frequencies.

**Online Resource 2.** Cross-amplification of 15 microsatellite markers developed for *Ruta lamarmorae* in two closely related species of the same genus. P, polymorphic locus; M, monomorphic locus; +, successful amplification but no information on whether the locus is polymorphic or not; -, failed amplification.

| Locus | *R. lamarmorae* | *R. corsica* | *R. chalepensis* |
| --- | --- | --- | --- |
| RL4 | P | P | P |
| RL5 | P | P | + |
| RL6 | P | P | - |
| RL7 | - | + | + |
| RL9 | P | P | + |
| RL10 | P | P | - |
| RL11 | P | P | P |
| RL12 | P | P | P |
| RL13 | P | P | + |
| RL15 | P | P | P |
| RL16 | P | P | + |
| RL17 | P | P | + |
| RL18 | P | P | + |
| RL19 | P | - | - |
| RL20 | - | - | - |

**Online Resource 3.** Pairwise population estimates of *R*_ST_. For abbreviations of populations see Table 1.

|  | *R. lamarmorae* |  | *R. corsica* |  |  |  |  |  |
| --- | --- | --- | --- | --- | --- | --- | --- | --- |
|  | BC | SS | MU | MC | SA | GH | AL | BA |
| BC |  |  |  |  |  |  |  |  |
| SS | 0.099 |  |  |  |  |  |  |  |
| MU | 0.093 | 0.031 |  |  |  |  |  |  |
| MC | 0.132 | 0.113 | 0.003 |  |  |  |  |  |
| SA | 0.253 | 0.147 | 0.080 | 0.160 |  |  |  |  |
| GH | 0.194 | 0.136 | 0.048 | 0.043 | 0.073 |  |  |  |
| AL | 0.184 | 0.052 | 0.046 | 0.057 | 0.191 | 0.059 |  |  |
| BA | 0.211 | 0.171 | 0.110 | 0.132 | 0.119 | 0.089 | 0.158 |  |

**Table S3.** Mean posterior probability of the data, Ln P(*X|K*), implemented by STRUCTURE over 20 runs for each *K* value. The model run here is an admixture model with correlated allele frequencies.

| $K$ | $\text{Ln } P(X\vert K)$<br>*R. corsica* | $\text{Ln } P(X\vert K)$<br>*R. lamarmorae* | $\text{Ln } P(X\vert K)$<br>*R. corsica + R. lamarmorae* |
| --- | --- | --- | --- |
| 1 | -3079.0 | -1968.5 | -5370.8 |
| 2 | -3050.0 | -1837.6 | -5081.6 |
| 3 | -2881.0 | -1808.7 | -4915.2 |
| 4 | -2859.0 |  | -4811.6 |
| 5 | -2906.2 |  | -4720.4 |
| 6 | -2929.2 |  | -4682.1 |
| 7 | -3023.8 |  | -4731.7 |
| 8 |  |  | -4807.3 |
| 9 |  |  | -4791.0 |

## Acknowledgements

M. M. and the project were funded by the Swiss National Science Foundation (SNSF) PMPDP3_129170. Participation to a conference was funded by the Claraz Schenkung.

